# Plant-derived CO_2_ mediates long-distance host location and quality assessment by a root herbivore

**DOI:** 10.1101/2020.03.12.988691

**Authors:** Carla C. M. Arce, Vanitha Theepan, Bernardus C. J. Schimmel, Geoffrey Jaffuel, Matthias Erb, Ricardo A. R. Machado

## Abstract

Insect herbivores can use volatile and visual cues to locate and select suitable host plants from a distance. The importance of CO_2_, arguable the most conserved volatile marker of metabolic activity, is not well understood in this context, even though many herbivores are known to respond to minute differences in CO_2_ concentrations. To address this gap of knowledge, we manipulated CO_2_ perception of the larvae of the western corn rootworm (*Diabrotica virgifera virgifera*; WCR) through RNA interference and studied how CO_2_ perception impacts their interaction with their host plant, maize (*Zea mays*). We show that the expression of a putative Group 2 carbon dioxide receptor, *DvvGr2*, is specifically required for dose-dependent larval responses to CO_2_ in the ppm range. Silencing *DvvGr2* has no effect on the ability of WCR larvae to locate host plants at short distance (<9 cm), but impairs host location at greater distances. Using soil arenas and olfactometer experiments in combination with *DvvGr2* silencing and CO_2_ scrubbing, we demonstrate that WCR larvae use CO_2_ as a long-range host plant finding cue, but employ other volatiles for short-range host location. We furthermore show that the larvae use CO_2_ as a fitness-relevant long-distance indicator of plant nutritional status: Maize plants that are well-fertilized emit more CO_2_ from their roots and are better hosts for WCR than plants that are nutrient-deficient, and the capacity of WCR larvae to distinguish between these plants depends exclusively on their capacity to perceive CO_2_ through *DvvGr2*. This study unravels how CO_2_ can mediate plant-herbivore interactions by serving as a distance-dependent host location and quality assessment cue.

## Introduction

Insect herbivores can use both visual and olfactory cues to locate suitable host plants from a distance. Volatile cues in particular can convey information about the identity and physiological status of a host plant and are integrated by herbivores to locate host plants for oviposition and feeding (***Visser and Avé, 1978***). Over the years, many attractive and repellent plant volatiles were identified (***Bruce et al., 2005; Späthe et al., 2013; Webster and Cardé, 2017***), and the importance of individual compounds and volatile blends was documented using synthetic chemicals (***Bruce and Pickett, 2011; Carrasco et al., 2015; Fraenkel, 1959; Natale et al., 2003; Visser and Avé, 1978***). More recently, molecular manipulative approaches were used to manipulate plant volatile production and herbivore perception *in vivo (**Fandino et al., 2019; Halitschke et al., 2008; Robert et al., 2013**)*, thus confirming the important role of plant volatiles in plant-herbivore interactions.

While the role of plant volatiles such as green-leaf volatiles, aromatic compounds and terpenes is well understood, much less is known about the role of plant-derived carbon dioxide (CO_2_) in plant-herbivore interactions. CO_2_ is a ubiquitous primary product of oxidative metabolism that is produced through the citric acid cycle (***Krebs and Johnson, 1937***), and is thus arguably the most conserved volatile marker of biological activity on the planet. Although plants are net sinks of CO_2_ due to their ability to fix carbon in the dark reactions of photosynthesis, they also release CO_2_ through respiration from metabolically active heterotrophic tissues such as flowers and roots, and autotrophic tissues such as leaves that operate below the light compensation point (e.g. at night). CO_2_ may thus be integrated into herbivore foraging activities as a marker of plant metabolic activity. *Datura wrightii* flowers for instance emit the highest levels of CO_2_ during times of high nectar availability; as hawkmoth pollinators are attracted to CO_2_, they may thus use this cue to locate rewarding flowers (***Goyret et al., 2008; Guerenstein et al., 2004; Guerenstein and Hildebrand, 2008; Stange, 1996; Stange, 1999; Stange and Stowe, 1999; Thom et al., 2004***). Similarly, lesions in apples result in high CO_2_ release and attract *Bactrocera tryoni* fruit flies. As CO_2_ at corresponding concentrations is attractive to the flies, it has been suggested that they may use plant-derived CO_2_ to locate suitable oviposition sites (**Stange, 1998**).

Root feeding insects are also highly attracted to CO_2_ *in vitro* (***Bernklau and Bjostad, 1998a, 1998b; Eilers et al., 2012; Hibbard and Bjostad, 1988; Jones and Coaker, 1978; Klingler, 1966; Nicolas and Sillans, 1989; Rogers et al., 2013; Strnad et al., 1986; Strnad and Dunn, 1990***). Given that CO_2_ is produced and released by plant roots and diffuses relatively well through the soil, a likely explanation for this phenomenon is that root herbivores use CO_2_ as a host location cue (***Bernklau and Bjostad, 1998a, 1998b; Doane et al., 1975; Erb et al., 2013; Johnson and Gregory, 2006; Johnson and Nielsen, 2012***), However, the reliability of CO_2_ as a host-location cue for root feeders has been questioned due to a number of reasons: i) CO_2_ can be emitted by many other sources apart from host plant roots, including decaying organic matter, microorganisms, and non-host plants, ii) there is a strong diurnal fluctuation in plant CO_2_ emissions that does not necessarily match with insect foraging habits and iii) other plant-released chemicals can be used by root herbivores for host location within a CO_2_ background (***Agus et al., 2010; Eilers et al., 2012; Erb et al., 2013; Hansen, 1977; Hibbard and Bjostad, 1988; Hiltpold and Turlings, 2012; Johnson and Nielsen, 2012; Reinecke et al., 2008; Weissteiner et al., 2012***). A model that may reconcile these different views is that CO_2_ may be used as an initial cue at long-distances, while other, more host specific volatiles may be used at shorter distances (***Erb et al., 2013; Johnson et al., 2006; Johnson and Nielsen, 2012***). So far, this model has not been experimentally validated, and the precise role of plant-derived CO_2_ as a host location cue by herbivores in general, and root herbivores in particular, remains unclear (***Eilers et al., 2016***). To the best of our knowledge, no studies so far have investigated the role of plant-derived CO_2_ in plant-herbivore interactions *in vivo* using molecular manipulative approaches.

The larvae of *Diabrotica virgifera virgifera* (western corn rootworm, WCR) feed exclusively on maize roots and cause major yield losses in the US and Eastern Europe (***Ciosi et al., 2008; Gray et al., 2009; Meinke et al., 2009; Toepfer et al., 2015***). The larvae rely on a number of volatile and non-volatile chemicals to identify and locate host plants, distinguish between suitable and less-suitable maize plants and forage within the maize root system (***Hiltpold et al., 2013; Johnson and Gregory, 2006; Johnson and Nielsen, 2012; Robert et al., 2012c; Schumann et al., 2018***). Non-volatile primary metabolites such as sugars and fatty acids as well as secondary metabolites such as benzoxazinoids and phenolic acid conjugates modulate larval behaviour (***Bernklau et al., 2011; Bernklau et al., 2015; Bernklau et al., 2016a; Bernklau et al., 2018b; Erb et al., 2015b; Hu et al., 2018; Huang et al., 2017; Robert et al., 2012c***). Volatiles including (*E*)-β-caryophyllene, ethylene and CO_2_ attract the larvae (***Bernklau and Bjostad, 1998a, 1998b; Robert et al., 2012b; Robert et al., 2012a***), while methyl anthranilate repels them (***Bernklau et al., 2016b***). Based on the finding that high CO_2_ levels can outweigh the attractive effects of other maize volatiles, it was suggested that CO_2_ may be the only relevant volatile attractant for WCR larvae (***Bernklau and Bjostad, 1998b***). However, under conditions where CO_2_ levels are similar, WCR larvae reliably choose between host plants of different suitability using other volatile cues (***Erb et al., 2011; Erb et al., 2015a; Huang et al., 2017; Lu et al., 2016; Robert et al., 2012b; Robert et al., 2012a***). The demonstrated ability of WCR larvae to respond to different volatile cues and the recent identification of putative CO_2_ receptors from transcriptomic data (***Rodrigues et al., 2016***) makes this species a suitable model system to investigate the role of CO_2_ in plant-herbivore interactions. Ongoing efforts to use CO_2_ as a bait to control WCR in the field (***Bernklau et al., 2004; Schumann et al., 2014b; Schumann et al., 2014a***) provide further motivation to assess the importance of this volatile for WCR foraging.

To understand the importance of CO_2_ for WCR foraging in the soil, we manipulated the insect’s capacity to perceive CO_2_. We reduced the expression levels of three putative WCR CO_2_ receptor-encoding genes through RNA interference (RNAi), resulting in the identification of *DvvGr2* as an essential gene for CO_2_ perception. Using *DvvGr2*-silenced larvae in combination with CO_2_-removal, we then assessed the importance of CO_2_ perception for WCR behaviour and foraging in olfactometers and soil arenas. Our experiments reveal how root-derived CO_2_ modulates the interaction between maize and its economically most damaging root pest and expand the current repertoire of adaptive explanations for the attraction of insect herbivores to CO_2_.

## Results

### The western corn rootworm (WCR) genome encodes three putative CO_2_ receptors

To identify genes encoding putative CO_2_ receptors in WCR, we used known CO_2_ receptor-encoding gene sequences as queries against the WCR genome (available from the National Center for Biotechnology Information (NCBI)). Three putative carbon dioxide receptor candidates, *DvvGr1, DvvGr2*, and *DvvGr3*, were identified, matching three candidate genes that were found in previous transcriptome analyses (Rodrigues *et al.*, 2016). Phylogenetic reconstruction based on *in silico* predicted protein sequences revealed orthologous relationships for the three WCR candidate receptors and the receptors of several other insects (Fig. 1A). Consistent with their taxonomy, we observed close homology between the protein sequences of the CO_2_ receptors of WCR and the protein sequences of other coleopteran insects such as *Tribolium castaneum* (Fig. 1A). Expression levels of *DvvGr1* and *DvvGr2* were found to be significantly higher in the head than in the rest of the body (torax and abdomen) of 2^nd^ instar WCR larvae (Figs. 1B, C). A similar trend was observed for *DvvGr3* (Fig. 1D). Protein tertiary structure and topology models indicated that all three genes encode for 7-transmembrane domain proteins, which is consistent with their roles as receptors (Fig. 1B-D).

**Figure 1.**
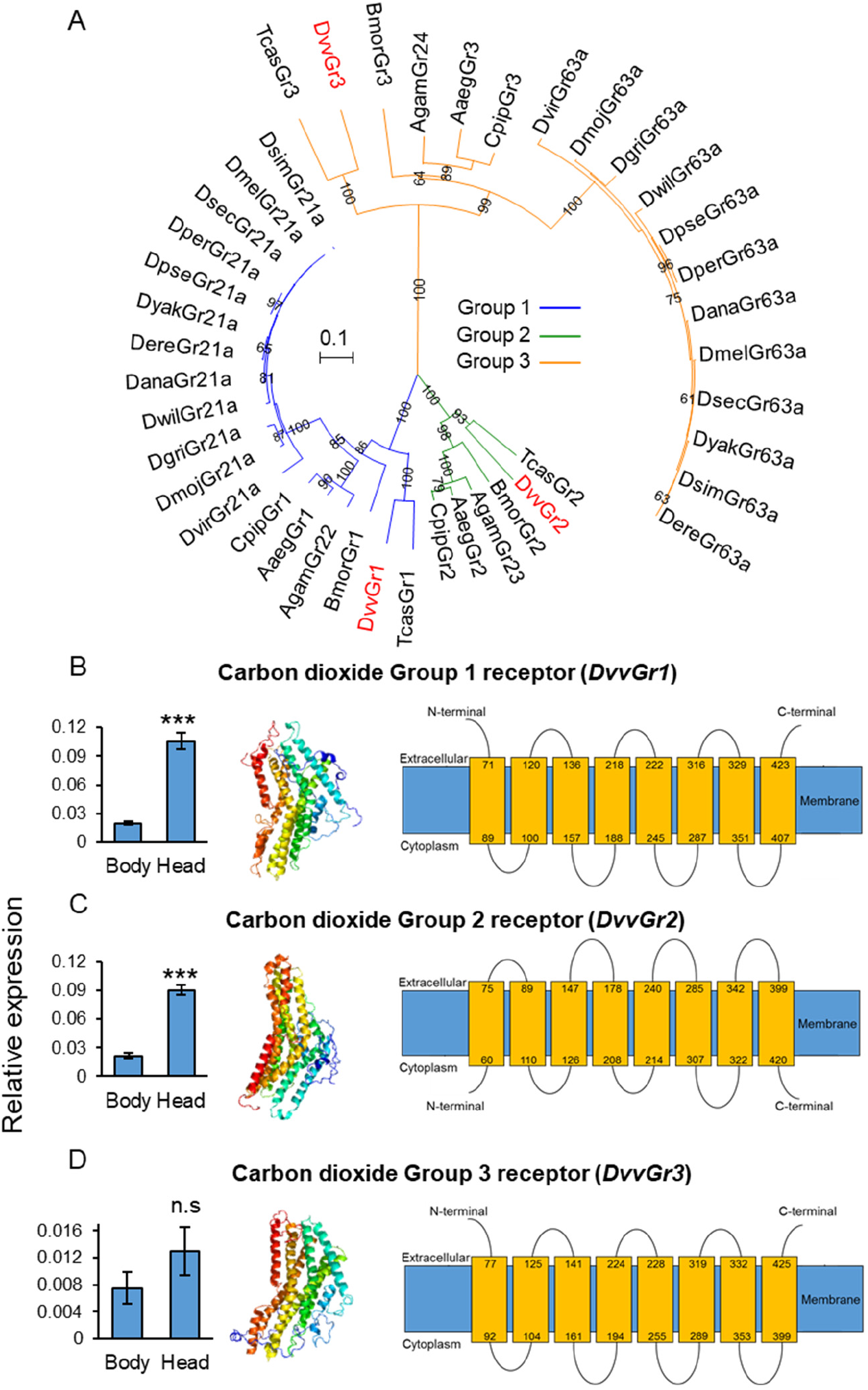
The WCR genome contains three putative carbon dioxide (CO_2_) receptors. (A) Phylogenetic relationships between putative CO_2_ receptors based on protein sequences of different insects. Dmel: *Drosophila melanogaster*, Dsim: *Drosophila simulans*, Dsec: *Drosophila sechellia*, Dyak: *Drosophila yakuba*, Dere: *Drosophila erecta*, Dana: *Drosophila ananassae*, Dper: *Drosophila persimilis*, Dpse: *Drosophila pseudoobscura*, Dwil: Drosophila willistoni, Dgri: *Drosophila grimshawi*, Dmoj: *Drosophila mojavensis*, Dvir: *Drosophila virilis*, Agam: *Aedes. gambiae*, Aaeg: *Aedes aegypti*, Cqui: *Culex quinquefasciatus*, Bmor: *Bombix mori*, Tcas *Tribolium castaneum*, Dvv*: Diabrotica virgifera virgifera* (Western corn rootworm, WCR). Mean (±SEM) relative gene expression levels of group 1 (*DvvGr1*) (B), group 2 (*DvvGr2*)(C), and group 3 (*DvvGr3*)(D) CO_2_ receptors in the bodies (thorax and abdomen) or heads of 2^nd^ instar WCR larvae (n=10). Asterisks indicate statistically significant differences between tissue types within genes (***: *p*<0.001 by one-way ANOVA with Holm’s multiple comparisions test; n.s.: not significant). (B-E) Predicted protein tertiary structure (left) and transmembrane protein topology (right) of: (B) *DvvGr1*; (C) *DvvGr2*; and (D) *DvvGr3* according to the Phyre2 algorithm.

### *DvvGr2* expression is specifically required for behavioural responses of WCR larvae to CO_2_

To determine the importance of *DvvGr1, DvvGr2*, and *DvvGr3* for the responsiveness of WCR larvae to CO_2_, we knocked down the expression of each gene individually through double-stranded RNA (dsRNA)-mediated RNA interference (RNAi) and conducted initial behavioural experiments with carbonated water as a CO_2_ source (Fig. 2). Oral administration of dsRNA targeting either *DvvGr1*, *DvvGr2*, or *DvvGr3* reduced the expression levels of these genes by 80%, 83% and 66% compared to WCR larvae fed with dsRNA of the green fluorescent protein (GFP) gene (herein referred as wild type, WT) (Fig. 2A). All RNAi constructs were confirmed to be gene specific (Fig 2B). Olfactometer arms containing carbonated water released approximately 100 ppm more CO_2_ than their counterparts filled with distilled water (Fig. 2B, Fig. S1A) and were more attractive to WT larvae than arms containing distilled water (Fig 2C). Silencing *DvvGr1* or *DvvGr3* expression did not alter this behaviour. In contrast, *DvvGr2*-silenced larvae did not show any preference any more (Fig 2C). To further explore the role of *DvvGr2* in WCR behaviour, we conducted a series of additional experiments. First, we assessed the impact of silencing *DvvGr2* on the capacity of WCR larvae to respond to other volatile and non-volatile host cues. *DvvGr2*-silenced larvae responded similarly to the repellent volatile methyl anthranilate (***Bernklau et al., 2016b; Bernklau et al., 2018a***) as WT larvae (Fig. 2D). Responsiveness to non-volatile attractants, Fe(III)(DIMBOA)_3_ and a blend of glucose, fructose and sucrose (***Bernklau et al., 2018b; Hu et al., 2018***) was also unaltered in *DvvGr2*-silenced larvae (Figs 2E, F), demonstrating that knocking down *DvvGr2* expression does not alter the capacity of WCR larvae to respond to other important chemical cues. Second, we assessed the contribution of *DvvGr2* to CO_2_ responsiveness using synthetic CO_2_ at different concentrations (Fig. 3, Fig. S1B). WT larvae showed characteristic dose-dependent behavioural responses to CO_2_. While they did not respond to 22 ppm CO_2_ above ambient CO_2_ levels, they were attracted to CO_2_ concentrations between 59-258 ppm above ambient, and repelled by CO_2_-enriched air at 950 ppm above ambient CO_2_ levels and above (Fig. 3). In contrast, *DvvGr2*-silenced larvae did not respond to CO_2_ enrichment at any of the tested concentrations (Fig. 3). These experiments show that WCR larvae are attracted to CO_2_ enriched environments within the physiological range of the maize rhizosphere (Fig. S1) and that *DvvGr2*-silencing fully and specifically suppresses CO_2_ responsiveness in WCR larvae.

**Figure 2.**
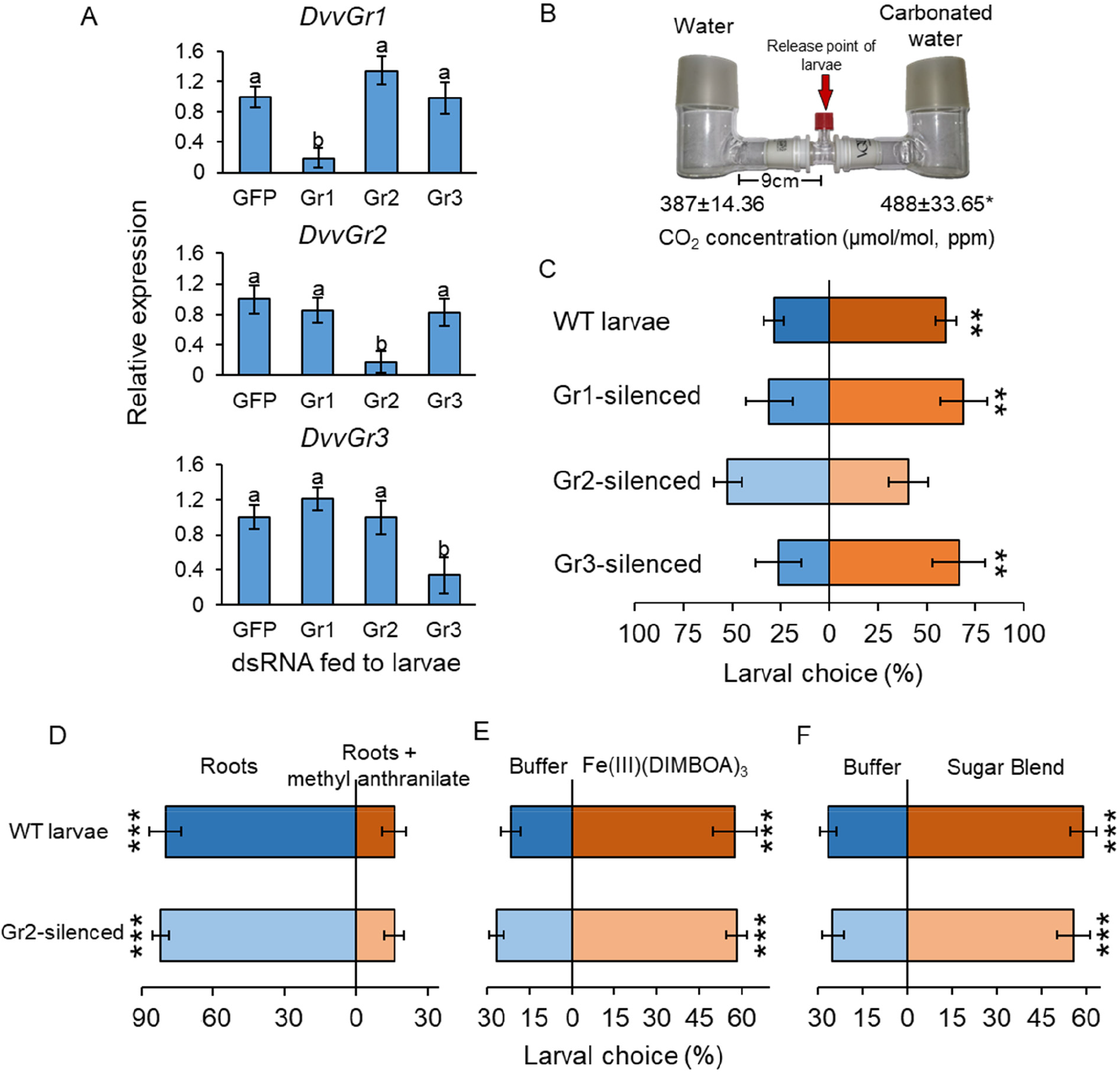
The carbon dioxide group 2 receptor (DvvGr2) is specifically required for behavioural responses of WCR to CO_2_. Mean (±SEM) relative gene expression levels of group 1 (*DvvGr1*), group 2 (*DvvGr2*), and group 3 (*DvvGr3*) CO_2_. receptors after WCR larvae were fed with dsRNA-expressing bacteria targeting green fluorescent protein (GFP, herein referred as WT), *DvvGr1*, *DvvGr2*, or *DvvGr3* genes (n=11-13). Different letters indicate significant differences of gene expression levels (*p*<0.05 by one-way ANOVA with Holm’s multiple comparisions test). (B) Mean (±SEM) CO_2_ levels in each arm of the two-arm belowground olfactometers used for behavioural experiments (n=4-8). Asterisk indicates significant differences in CO_2_ levels (*: *p*<0.05 by one-way ANOVA with Holm’s multiple-comparisons test). Mean (±SEM) proportion of WCR larvae observed in the olfactometer arms with higher CO_2_ levels (carbonated water side) or in control arms (distilled water side). Seven olfactometers with six larvae each were assayed (n=7). Mean (±SEM) proportion of WCR larvae observed on roots or on roots placed next to filter paper discs impregnated with methyl anthranilate (D), on filter paper discs impregnated with buffer or with Fe(III)(DIMBOA)_3_ (E), and on filter paper discs impregnated with buffer or with a blend of glucose, fructose, and sucrose (F). For panel D, ten choice arenas with five larvae each were assayed (n=10). For panels E and F, twenty choice arenas with six larvae each were assayed (n=20). Asterisks indicate statistically significant differences in larval choices between treatments (***: *p*<0.001 by two-way ANOVA followed by FDR-corrected post hoc tests). For *P*-values of Generalized Linear Models (GLMs), refer to Table S1. Refer to Fig. S1A for quantities of CO_2_ in each arm.

**Figure 3.**
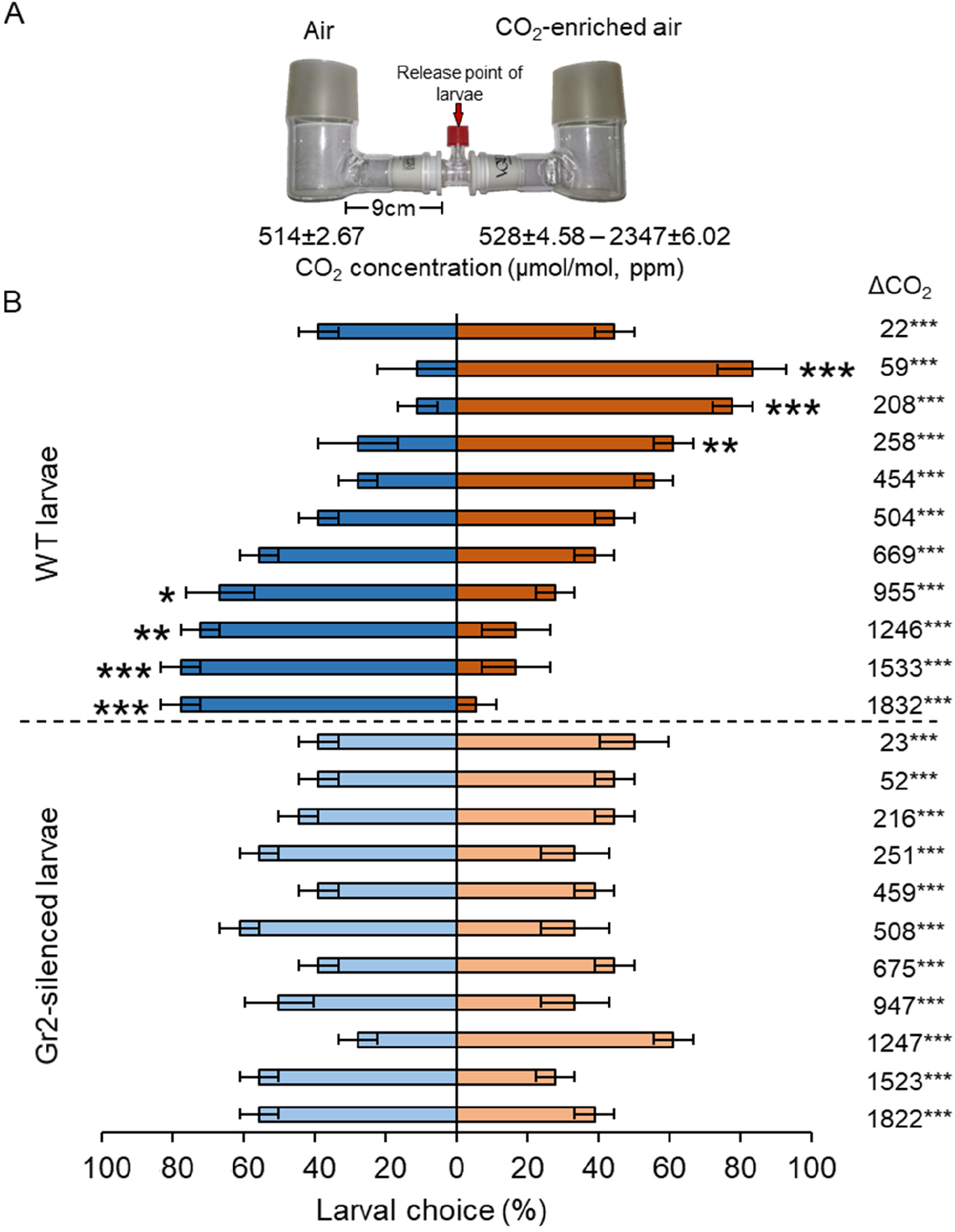
DvvGr2 is required for dose-dependent behavioural responses of WCR to CO_2_. (A) Olfactometer setup used for behavioural experiments. (A) Mean (±SEM) CO_2_ levels in each arm of the olfactometers (n=3). (B) Mean (±SEM) proportion of WCR larvae observed in each arm of the olfactometers. Three olfactometers with six larvae each were assayed (n=3). Asterisks indicate statistically significant differences between larval choices (*: *p*<0.05; **: *p*<0.01; ***: *p*<0.001 by three-way ANOVA followed by FDR-corrected post hoc tests). For *P*-values of Generalized Linear Models (GLMs), refer to Table S1. ΔCO_2_ values are given for each level. Asterisks indicate significant differences between CO_2_ levels in the two olfactometer arms (***: p<0.001 by three-way ANOVA followed by FDR-corrected post hoc tests). Refer to Fig. S1B for quantities of CO_2_ in each arm.

### *DvvGr2* expression does not impair larval motility or short-range host location

To assess the impact of *DvvGr2* on larval motility, we followed the trajectories of individual larvae in Petri plates that were outfitted with a CO_2_ point releaser (Fig. 4A-C). WT larvae made frequent turns, but consistently oriented themselves towards the CO_2_ release point. Once they reached the CO_2_ release point, they stopped moving (Fig. 4A). *DvvGr2*-silenced larvae exhibited similar turning behaviour as WT larvae, but did not move towards the CO_2_ release point (Fig. 4B). WT larvae spend more time on CO_2_ release point than *DvvGr2*-silenced larvae (Fig. 4C). During the movement phase, the mean speed of WT larvae and *DvvGr2*-silenced larvae was similar (Fig. 4C), but the distance covered by *DvvGr2*-silenced larvae was higher, as they did not stop at the CO_2_ release point. In a second experiment, we followed the trajectories of individual larvae in Petri plates with maize roots (Fig. 4D, E). Speed and distance covered were similar between WT and *DvvGr2*-silenced larvae (Fig. 4F). Surprisingly, both WT and *DvvGr2*-silenced larvae oriented themselves towards the maize roots and reached the maize roots after a similar amount of time (Fig. 4F). This result shows that *DvvGr2* expression is required for the location and detection of CO_2_, but does not influence WCR motility nor its ability to locate maize roots over short distances (i.e.: <9 cm).

**Figure 4.**
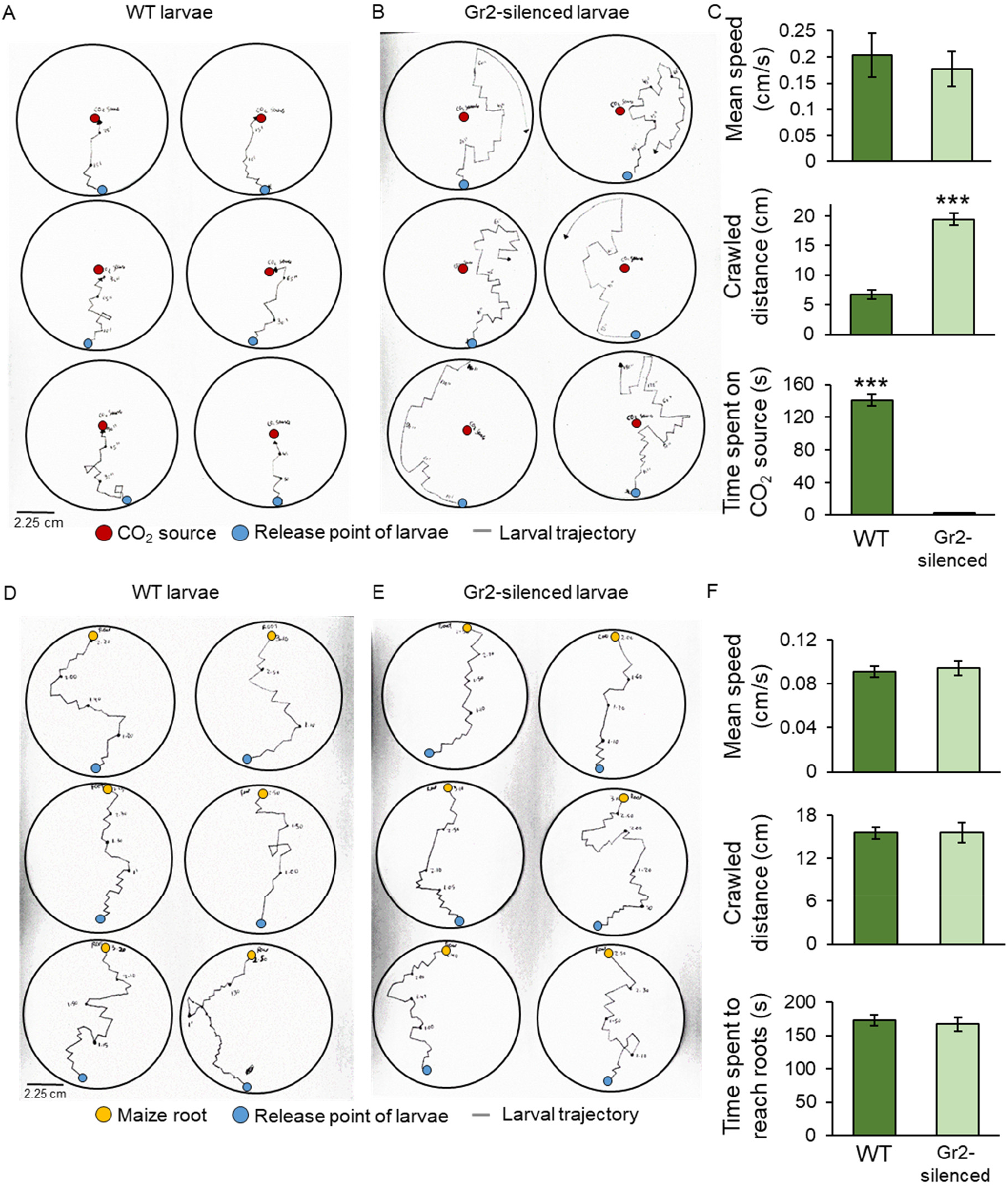
Silencing the carbon dioxide group 2 receptor (DvvGr2) impairs WCR responses to CO_2_ without changing motility or search behavior. (A-B) Trajectories of individual WT (A) and *DvvGr2*-silenced (B) WCR larvae in Petri plates with a CO_2_ source. The blue circles represent larval release points. The red circles represent CO_2_ sources consisting of a fine needle that releases CO_2_ at 60 ppm above ambient. (C) Mean (±SEM) speed and distance covered during the movement phase, and time spent at the CO_2_ source during the first 3 min of the experiment. (D-E) Trajectories followed by WT (D) and by *DvvGr2*-silenced (E) WCR larvae on Petri plates containing maize seedling roots. The blue circles represent larval release points. The yellow circles represent maize seedling roots. (F) Mean (±SEM) speed and distance covered during the movement phase, and time necessary to reach the maize root during the first 3 min of the experiment. (F). For both experiments, six Petri plates with one larva each were assayed (n=6). Asterisks indicate statistically significant differences between mobility parameters of WT and *DvvGr2*-silenced larvae (***: *p*<0.001 by one-way ANOVA followed by Holm’s multiple-comparisons test).

### *DvvGr2*-dependent CO_2_ responsiveness mediates long-range host location

To further explore the role of CO_2_ and *DvvGr2* in volatile-mediated host location, we performed a series of olfactometer experiments with maize plants grown in sand on one side and sand only on the other side. We varied the distance between the volatile sources and the release points of the larvae between 9 and 18 cm (Fig. 5). We also manipulated CO_2_ emission of maize roots in a subset of olfactometers by adding a layer of CO_2_-absorbing soda lime into the olfactometer arms. CO_2_ measurements revealed that the presence of a host root system increased CO_2_ concentrations by approx. 100 ppm above ambient CO_2_ levels in the corresponding olfactometer arm (Fig. 5, Fig. S1C). The soda lime reduced ambient CO_2_ concentrations in the olfactometer arms by approx. 100 ppm, and equalized CO_2_ concentrations between arms with and without a host plant (Fig. 5), thus validating the CO_2_ scrubbing approach. Larvae did not have direct access to the plant, the plant growth medium or the soda lime, and received no visual cues, and thus had to rely on host plant volatiles for orientation. When released at distance of 9 cm from the volatile sources, both WT and *DvvGr2*-silenced larvae showed a clear preference for the olfactometer arms leading to host plants (Fig. 5A). This preference was still intact in olfactometers outfitted with a soda lime filters, confirming that volatiles other than CO_2_ are sufficient for volatile-mediated host location at a short distance. At a distance of 18 cm from the volatile sources, WT larvae showed a similarly strong preference for arms leading to host plants (Fig. 5B). By contrast, *DvvGr2*-silenced larvae did not exhibit any preference (Fig. 5B). In the presence of a soda lime filter, neither WT nor *DvvGr2*-silenced larvae were attracted to arms with a host plant (Fig. 5B). Taken together, these experiments demonstrate that WCR larvae require plant-derived CO_2_ to locate host plants over longer distances in a *DvvGr2*-dependent manner.

**Figure 5.**
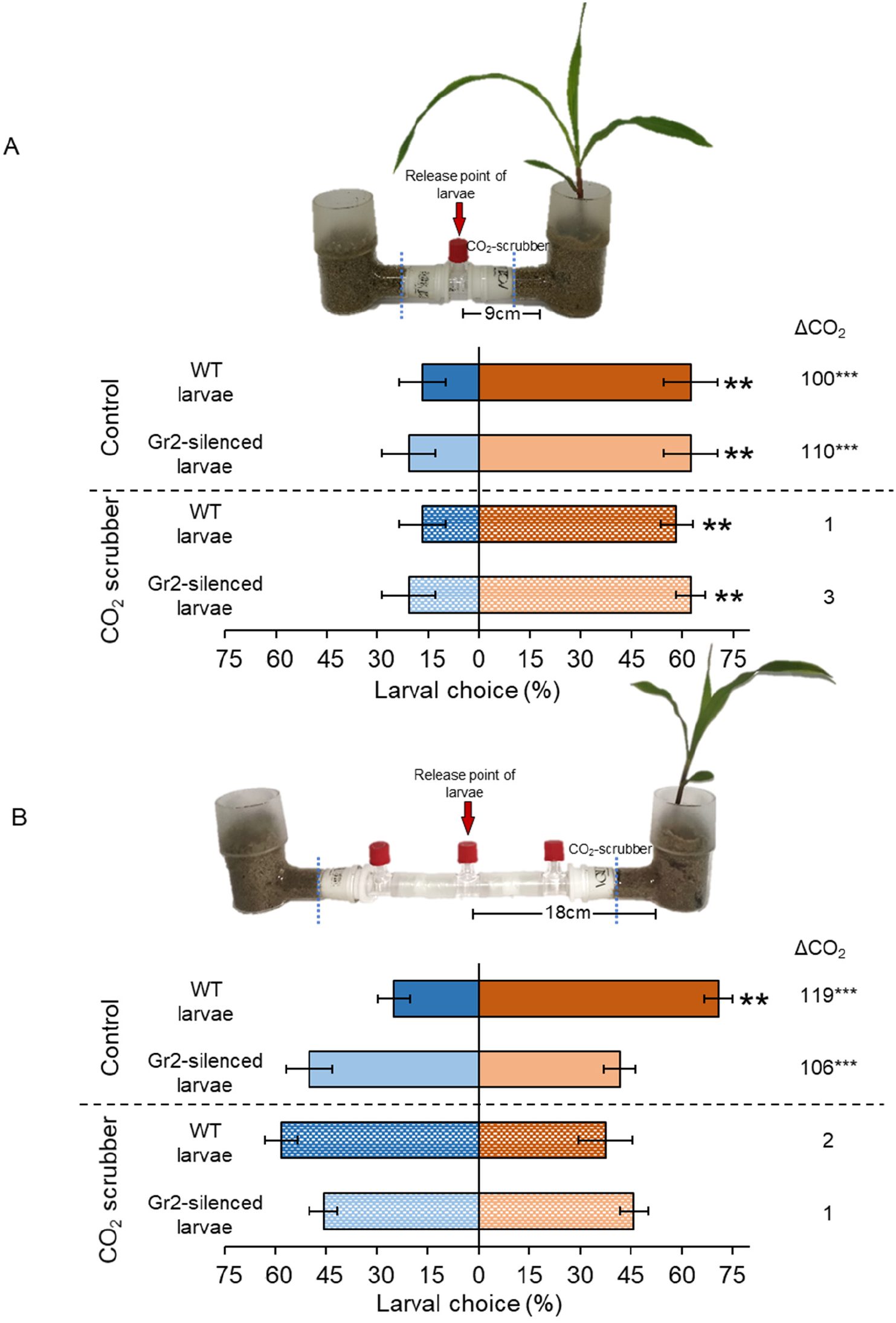
Plant-derived CO_2_ mediates DvvGr2-dependent long-distance host location. (A, B) Choice (mean±SEM) of WCR larvae in two-arm olfactometers. Control olfactometers allowed for plant-derived CO_2_ to diffuse into the choice chamber, while CO_2_ scrubber olfactometers were outfitted with soda lime to suppress CO_2_ diffusion. ΔCO_2_ represents the difference in CO_2_ in the arms with plants compared to arms without plants (***p<0.001). For detailed data on CO_2_ levels, refer to Fig. S1C. Olfactometer arms were either 9 cm (A) or 18 cm (B) in length. Four olfactometers with six larvae each were assayed using WT or *DvvGr2*-silenced larvae (n=4). Asterisks indicate statistically significant differences between treatments (**: *p*<0.01 by four-way ANOVA followed by FDR-corrected post hoc tests). For *P*-values of Generalized Linear Models (GLMs), refer to Table S1.

### Root-derived CO_2_ enhances volatile-mediated host location over longer distances

WCR larvae can move up to 1 m in the soil. Second and third instar larvae in particular are known to move between maize plants across rows in maize fields (***Hibbard et al., 2003***). To test whether *DvvGr2*-mediated CO_2_ responsiveness mediates host location over longer distances in a soil context, we planted maize plants in soil-filled plastic trays, released WCR larvae at distances of 16, 32, 48 or 64 cm from the maize plants and evaluated larval positions after 8 h (Fig. 6). This time point was chosen based on preliminary observations showing that larvae take approx. 8 h to cross the soil arenas. The CO_2_ emitted by maize roots formed a gradient in the soil, starting at about 506 ppm in the rhizosphere (Zone 1), and 430 ppm at distances of 16-32 cm from the plant (Zone 2) (Fig. 6B, Fig. S1F). At distances of more than 32 cm from the plant, the CO_2_ levels were around 400 ppm and statistically indistinguishable from soil without plants or ambient air (Fig. S1F). To confirm that larval motility is not altered by *DvvGr2*-silencing in a soil context, we first released WT and *DvvGr2*-silenced larvae into the middle of a set of arenas without a host plant and evaluated larval positions after 8 h. We found that the larvae dispersed equally across the arenas, without any difference between WT and *DvvGr2*-silenced larvae (Fig. 6A). Eight hours after releasing the larvae into arenas that included host plants on one side, 53% of WT larvae that were released at 64 cm from the plant were retrieved from the rhizosphere, i.e.: in Zone 1 (Fig. 6C). In contrast, only 33% of the *DvvGr2*-silenced larvae that were released at the same distance were recovered from the maize rhizosphere (Fig. 6C). Significantly more *DvvGr2*-silenced larvae were recovered further away from the plants, in zones 3 and 4 (Fig 6C). The number of WT and *DvvGr2*-silenced WCR larvae found close to the host plant increased with decreasing release distance, as did the difference between WT and *DvvGr2*-silenced larvae (Fig. 6C-F). At a release distance of 16 cm, only slightly more WT than *DvvGr2*-silenced larvae were found close to the plant roots (Fig 6F). To further confirm the role of *DvvGr2* in mediating host plant location over long distances in the soil, we performed a time-course experiment where we released WT and *DvvGr2*-silenced larvae in Zone 5 (64 cm away from the host plant) and then recorded how rapidly they reached Zone 1 containing host plants (Fig. S2). The capacity of the larvae to directly feed on the host roots was impeded using a volatile-permeable root barrier (Fig. S2A). Within 10 h, 36% of the released WT larvae were found in Zone 1, and within 32 h, this number had increased to 90% (Fig. S2B). By contrast, only 24% of the released *DvvGr2*-silenced larvae were found in Zone 1 after 10 h, and after 32 h, this value had only increased to 56% (Fig. S2B). Thus, the capacity to detect CO_2_ gradients contributes to successful host location by WCR larvae in a distance specific manner in the soil.

**Figure 6.**
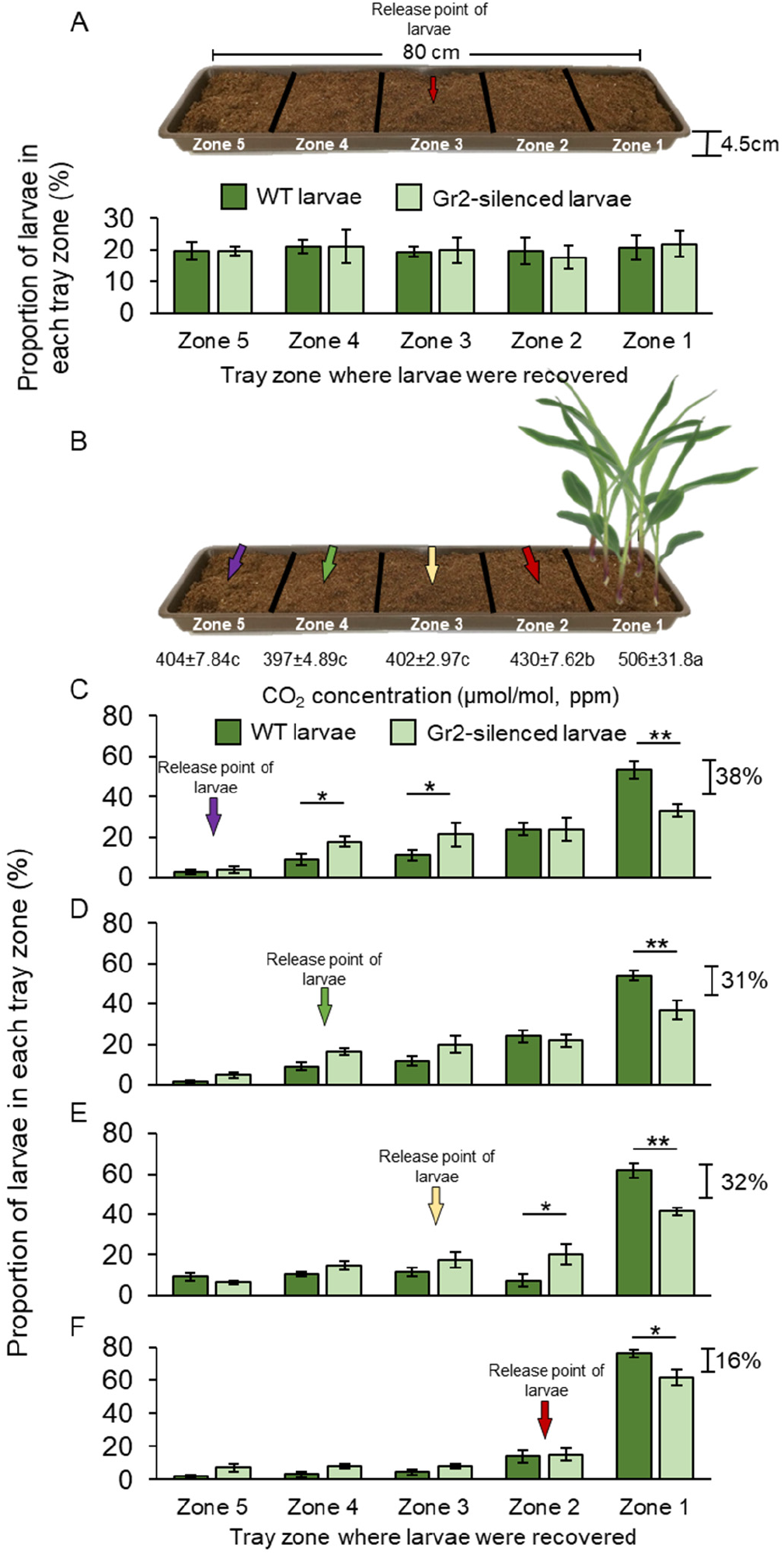
Root-derived CO_2_ enhances WCR host location over long distances in the soil. (A) Mean (±SEM) proportion of WT (dark green) or *DvvGr2*-silenced (light green) WCR larvae observed in the different tray zones eight hours after releasing the larvae in the centre of soil-filled trays without plants. Three trays per larval type with twenty larvae each were assayed (n=3). Schematic representation (photomontage) of experimental setup used for behavioural experiments depicting mean (±SEM) CO_2_ levels detected in the soil gas phase of each tray zone (n=3-4). Different letters indicate significant differences in CO_2_ levels (*p*<0.05 by one-way ANOVA with Holm’s multiple-comparisons test). (C-F) Mean (±SEM) proportion of WT (dark green) or *DvvGr2*-silenced (light green) WCR larvae observed in the different tray zones eight hours after releasing the larvae at distances of 64 cm (C), 48 cm (D), 32 cm (E), or 16 cm (F) from the plants. Six trays per larval type and distance combination with twenty larvae each were assayed (n=6). Asterisks indicate statistically significant differences in the proportion of WT and *DvvGr2*-silenced larvae found in each tray zone (*: *p*<0.05; **: *p*<0.01 by three-way ANOVA followed by FDR-corrected post hoc tests). For *P*-values of Generalized Linear Models (GLMs), refer to Table S1. For detailed data on CO_2_ levels, refer to Fig. S1E.

### Root-derived CO_2_ is used for host quality assessment by WCR larvae

Plant nutritional status determines plant growth and defense, and can thus modulate plant-herbivore interactions (***Wetzel et al., 2016***). To test for a possible connection between plant nutritional status, host suitability and CO_2_-dependent herbivore attraction, we varied the nutrient supply of maize plants and then carried out CO_2_ measurements, and behavioural and insect performance experiments (Fig. 7). Plants that were well fertilized released higher levels of CO_2_ into the soil than did plants that received a medium (50 % of optimally fertilized plants) or low (10% of optimally fertilized plants) dose of fertilizer (Fig 7A, B). As observed before, soil CO_2_ levels decreased with increasing distance from the plants and were lowest in the middle of the experimental trays (Fig. 7A, B). In choice experiments with maize plants planted approx. 50 cm apart, which corresponds to row spacing used for high planting densities in maize cultivation, WT larvae showed a significant preference for well-fertilized over medium-or low-fertilized plants (Fig 7C, D). *DvvGr2*-silenced larvae did not distinguish between plants subjected to the different fertilizer regimes (Fig. 7C, D). In no-choice experiments, WCR larvae gained most weight on well-fertilized maize plants than on plants treated with medium or low doses of fertilizer (Fig. 7E). Hence, intact CO_2_ perception is required for WCR larvae to express host quality related preferences for maize plants growing at agriculturally relevant distances.

**Figure 7.**
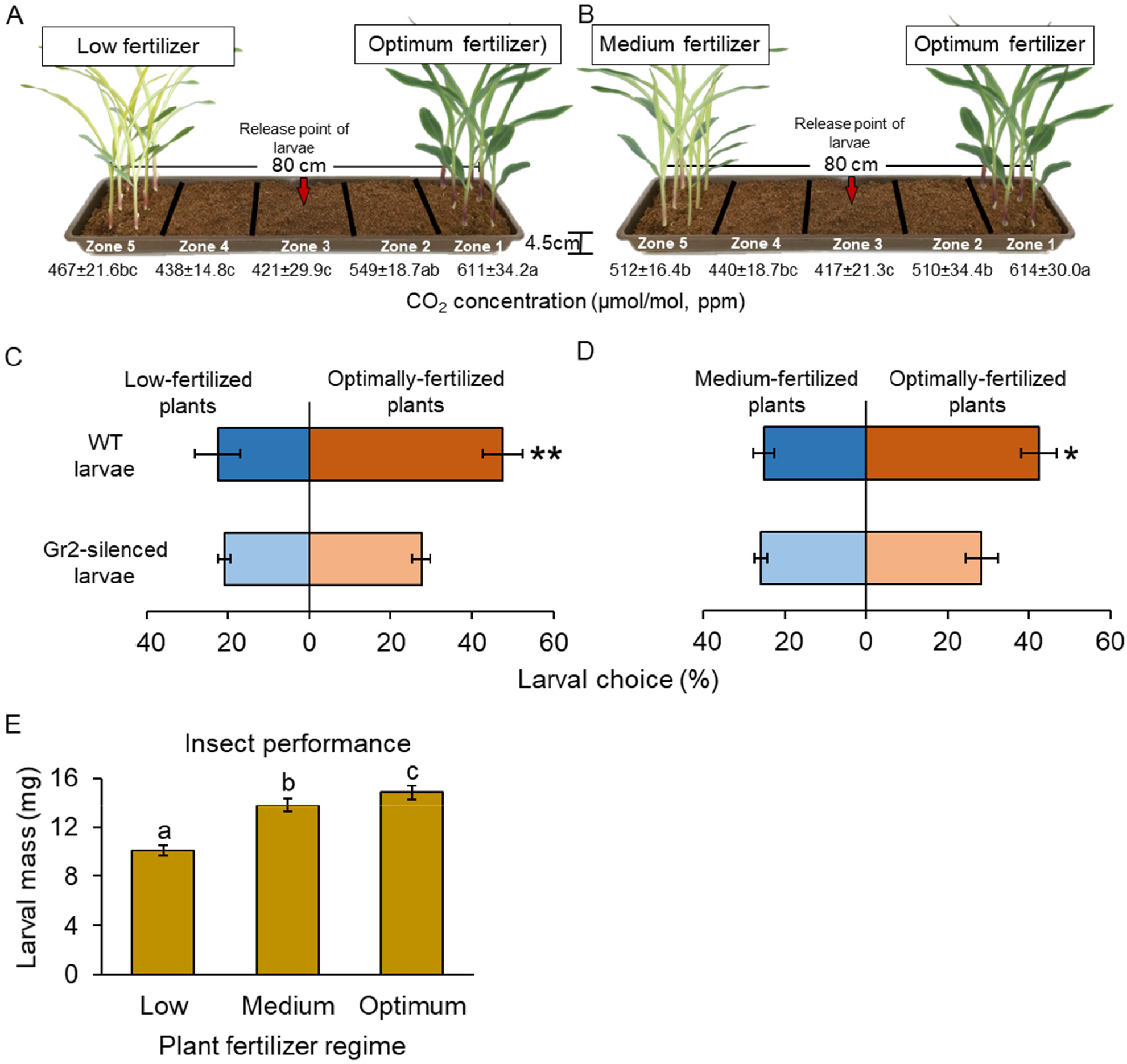
Root-derived CO_2_ is used for host quality assessment by WCR larvae in the soil. Schematic representation (photomontage) of experimental set up used to evaluate insect preference for plants fertilized with low or optimum fertilizer doses (A), and for plants fertilized with medium or optimum fertilizer doses (B) depicting mean (±SEM) CO_2_ levels detected in the soil gas phase of each tray zone (n=10). Different letters indicate statistically significant differences in CO_2_ levels (*p*<0.05 by one-way ANOVA with Holm’s multiple comparisions test). (C, D) Mean (±SEM) proportion of WCR larvae recovered from the rhizosphere (either Zone 1, or Zone 5) in choice assays contrasting larval preference for plants fertilized with low or optimum fertilizer doses (C), and for plants fertilized with medium or optimum fertilizer doses (D) eight hours after releasing the larvae. Six trays with twenty larvae per tray were assayed (n=6). Different letters indicate statistically significant differences in larval preferences (*: *p*<0.05; **: (*: *p*<0.01 by two-way ANOVA followed by FDR-corrected post hoc tests). (E) Mean (±SEM) weight of WCR larvae after 7d feeding on plants fertilized with low, medium or optimum fertilizer doses. Twenty solo cups with 4 to 7 larvae each were assayed (n=20). Different letters indicate statistically significant differences in larval mass (*p*<0.05 by one-way ANOVA followed by Holm’s multiple comparisions tests). For *P*-values of Generalized Linear Models (GLMs), refer to Table S1. For detailed data on CO_2_ levels, refer to Fig. S1D.

## Discussion

Volatile chemicals have a strong impact on the behaviour and performance of organisms across the tree of life (***Arce et al., 2017; Bentley and Day, 1989; Gang and Hallem, 2016; Hansson and Stensmyr, 2011; Johnson and Gregory, 2006; Miller and Strickler, 1984; Veyrat et al., 2016; Webster and Cardé, 2017; Ye et al., 2018***). Many organisms, including bacteria, fungi, nematodes, plants, insects, fish and mammals respond to CO_2_, for instance (***Bahn and Mühlschlegel, 2006; Cummins et al., 2014; Eilers et al., 2012; Hallem and Sternberg, 2008; Hester et al., 2012; Huckstepp and Dale, 2011; Jones et al., 2007; Perry and Abdallah, 2012; Raguso et al., 2005; Stange and Stowe, 1999***). However, the role of CO_2_ in plant-herbivore interactions is poorly understood and has not been studied *in vivo* through molecular manipulation approaches.

In this study, we conducted gene sequence similarity analyses, phylogenetic relationship reconstructions, RNA interference and behavioural experiments with different setups to explore the biological relevance of root-derived CO_2_ for plant-herbivore interactions. We found that the WCR genome contains at least three putative CO_2_ receptor-encoding genes: *DvvGr1*, *DvvGr2*, and *DvvGr3*, which is consistent with previous transcriptomic-based studies (***Rodrigues et al., 2016***). Protein tertiary structure and topology prediction models show that the identified genes code for proteins that contain seven transmembrane domains, which is consistent with the protein topology of gustatory and odor receptors (***Dahanukar et al., 2005; Hallem et al., 2006***). Larval behaviour and gene silencing based-functional characterization of the three identified WCR putative CO_2_ receptor genes revealed that the intact expression of *DvvGr2*, is essential for the attractive effects of CO_2_ to WCR larvae. Knocking down *DvvGr2* rendered larvae fully unresponsive to synthetic and plant-derived CO_2_. In *Aedes aegypti, Helicoverpa armigera* and *Drosophila melanoganster*, both carbon dioxide receptors *Gr1* and *Gr3* are required for CO_2_ detection (***Erdelyan et al., 2012; Jones et al., 2007; Kwon et al., 2007; McMeniman et al., 2014; Ning et al., 2016; Suh et al., 2004***). In *Culex quinquefasciatus*, both *Gr2* and *Gr3* carbon dioxide receptors are required, while *Gr1* act as a modulator (***Xu et al., 2020***). In *Aedes aegypti*, the involvement of *Gr2* in carbon dioxide responsiveness is still under debate (***Erdelyan et al., 2012; Kumar et al., 2019***). Taken together, the molecular elements required for carbon dioxide perception may be species-specific. Our results support this notion, as *DvvGr2*, but not *DvvGr1* and *DvvGr3*, are crucial for CO_2_ responsiveness. The role of *DvvGr1* and *DvvGr3* for WCR remains to be determined, but their presence and expression may hint at additional complexity in developmental and/or tissue-specific patterns of CO_2_ responsiveness in this species.

Despite the inability of *DvvGr2*-silenced WCR larvae to respond to differences in CO_2_ levels, the larvae were still able to orient towards maize roots at short distances of 8-10 cm. Olfactometer experiments in combination with a CO_2_ scrubber that removes plant-derived CO_2_ demonstrates that other volatile cues can be used by WCR larvae to locate maize plants at short distances. Earlier studies found that (*E*)-β-caryophyllene, which is emitted from the roots of certain maize genotypes when they are attacked by root herbivores, attracts second and third instar WCR larvae and allows them to aggregate on maize plants and thereby enhance their fitness (***Robert et al., 2012b***), while neonate larvae are not attracted to this volatile (***Hiltpold and Hibbard, 2016***). Ethylene has also been shown to attract WCR larvae (***Robert et al., 2012a***), and MBOA or its breakdown products have also been proposed as volatile attractants (***Bjostad and Hibbard, 1992***). Methyl anthranilate on the other hand has been shown to repel WCR larvae (***Bernklau et al., 2016b; Bernklau et al., 2018a***). Many other leaf- and root feeding herbivores are known to respond to plant volatiles other than CO_2_ (***Bruce et al., 2005***). Given the low reliability of CO_2_ as a host-specific cue, it is probably not surprising that WCR, as a highly specialized maize feeder, can use other volatile cues to locate host plants. Integrating other volatile cues likely allows WCR larvae to locate maize plants even in the absence of reliable CO_2_ gradients in the soil, thus increasing the robustness of its foraging behaviour at short distances. An intriguing result in this context is the fact that WCR larvae show the same efficiency in locating maize roots at short distances in the absence of a CO_2_ gradient, suggesting that this volatile may not play a role as a cue at close range.

Although intact CO_2_ perception was not required for host location at short distances, it had a strong impact on the capacity of WCR larvae to reach the maize rhizosphere at long distances. The advantages of CO_2_ are that it is abundantly produced through respiration by most organisms, that it is relatively stable (***Jones and Coaker, 1977; Li et al., 2016***), and that it diffuses rapidly in air, water and soil (***Hashimoto and Suzuki, 2002; Ma et al., 2013***). CO_2_ may thus be a suitable long-range cue to locate organisms with high respiratory rates, such as mammals and heterotroph plant parts, including roots (***Johnson and Nielsen, 2012***). Aboveground insects can be attracted to CO_2_ traps located as far away as 10 m, and it is estimated that this distance could even be as long as 60 m under optimal environmental circumstances (***Guerenstein and Hildebrand, 2008; Zollner et al., 2004***). For belowground insects, this distance is hypothesized to be within the lower cm-range as CO_2_ diffusion is substantially decreased within the soil matrix compared to the air (***Bernklau et al., 2005; Doane et al., 1975; Doane and Klingler, 1978; Klingler, 1966***). Other volatiles that are less abundant and diffuse even less well through the soil such as (*E*)-β-caryophyllene are unlikely to be detectable at distances of more than 10 cm (***Chiriboga et al., 2017; Hiltpold and Turlings, 2008***). In support of this, we did not succeed in the past to detect maize volatiles in the soil at distance of more than 10 cm, even using a highly sensitive combination of solid-phase microextraction (SPME) and gas chromatography/mass spectrometry (GC/MS). These volatiles are thus likely useful host location cues at short, but not long distances in the soil. The finding that WCR integrates CO_2_ perception with other environmental cues and that attraction to CO_2_ is context dependent is in line with patterns reported for other insects such as mosquitoes, whose response to stimuli such as color, temperature and human body odors is enhanced by CO_2_ (***McMeniman et al., 2014; van Breugel et al., 2015***), and pollinating hawkmoths, that use CO_2_ as a redundant volatile distance stimulus in a sex-specific manner (***Goyret et al., 2008***).

A recent study shows that a CO_2_ receptor in *Drosophila* flies is also involved in the detection and behavioral responses to other volatiles (***MacWilliam et al., 2018***). We observed that *DvvGr2*-silenced larvae were repelled by methyl anthranilate, a potent maize root repellent, to a similar extent than WT larvae, suggesting that their sensitivity to other plant volatiles is unchanged (***Bernklau et al., 2016b***). In *Drosophila* flies, the CO_2_ receptor *Gr63a* is required for spermidine attractiveness over short time spans, i.e.: less than 1 min, but not over longer time spans (hours), when other receptors likely become more important (***MacWilliam et al., 2018***). In the present experiments, WCR behaviour was evaluated after one or more hours. The CO_2_ scrubber experiment provides further evidence that the foraging patterns observed in this study are not due to different sensitivity of *DvvGr2*-silenced larvae to other root volatiles.

Apart from acting as a long-distance host location cue, CO_2_ also links plant fertilization to herbivore behaviour by guiding WCR to well fertilized plants. As WCR larvae are resistant to root defenses of maize (***Robert et al., 2012b***), it is likely to benefit from increased fertilization, independently of the plant’s defensive status. As the plant nutritional status and host quality for WCR larvae is associated with higher CO_2_ release from the roots, following the highest concentrations of CO_2_ in the soil may be adaptive for the herbivore, as it may increase its chance not only to find a maize plant *per se*, but also to identify a plant that has the resources to grow vigorously and that is a better host. WCR larvae are specialized maize pests that have evolved with intense maize cultivation in the corn belt of the US (***Gray et al., 2009***) and are resistant to maize defense metabolites (***Robert et al., 2012b***). Following the strongest CO_2_ gradient in an equally spaced maize monoculture may indeed be a useful strategy for this root feeder to locate suitable food sources. An association between CO_2_ emission and food-source profitability was also suggested for *Datura* flowers, that emit the highest level of CO_2_ in times when nectar is most abundant (***Guerenstein et al., 2004; Thom et al., 2004***). These findings support the hypothesis that CO_2_ is used as a marker of metabolic activity that allows for an assessment of the vigour and profitability of a wide variety of hosts.

In summary, this work demonstrates how an herbivore uses its capacity to perceive CO_2_ to locate host plants and to distinguish plants of different quality. Volatiles other than CO_2_ are also integrated into host finding behaviour in the soil, but their effects are more important at shorter distances. Thus, evidence is now accumulating that CO_2_ acts as a fitness-relevant host location cue in different insects, likely because of its unique role as a highly conserved long-range marker of metabolic activity within complex sensory landscapes.

## Materials and Methods

### Plants and planting conditions

Maize seeds (*Zea mays* L., var. Akku) were provided by Delley Semences et Plantes SA (Delley, CHE). Seedlings were grown under greenhouse conditions (23 ± 2°C, 60% relative humidity, 16:8 h L/D, and 250 mmol/m^2^/s^1^ additional light supplied by sodium lamps). Plantaaktiv^®^ 16+6+26 Typ K fertilizer (Hauert HBG Dünger AG, Grossaffoltern, Switzerland) was added twice a week after plant emergence following the manufacturer’s recommendations. The composition of the fertilizer is: total nitrogen (N) 16%, nitrate 11%, nmmonium 5%, phosphate (P_2_O_5_) 6%, potassium oxide (K_2_O) 26%, magnesium oxide (MgO) 3.3%, borum (B) 0.02%, cooper (Cu, EDTA-chelated) 0.04%, iron (Fe, EDTA-chelated) 0.1%, manganese (Mn, EDTA-chelated) 0.05%, molybdenum (Mo) 0.01%, and zink (Zn, EDTA-chelated) 0.01%. When plants were used as insect food, seedlings were germinated in vermiculite (particle size: 2-4 mm; tabaksamen, Switzerland) and used within 4 days after germination.

### Insects and insect rearing

*Diabrotica virgifera virgifera* (Western corn rootworm, WCR) insects used in this study were derived from a non-diapausing colony reared at the University of Neuchâtel. The eggs used to establish the colony were originally supplied by USDA-ARS-NCARL, Brookings, SD. Upon hatching, insects were maintained in organic soil (Selmaterra, Bigler Samen AG, Thun, Switzerland) and fed freshly germinated maize seedlings (var. Akku).

### Identification of CO_2_ receptor genes

To identify CO_2_ receptor orthologues in WCR, we used CO_2_ receptor-encoding gene sequences of *Tribolium castaneum* and several sequences from other insects as queries against publicly available WCR genome sequences (NCBI accession: PXJM00000000.2) using the National Center for Biotechnology Information Basic Local Alignment Search Tool (NCBI BLAST)(***Robertson and Kent, 2009; Wang et al., 2013; Xu and Anderson, 2015***). The full gene sequences can be retrieved from the National Center for Biotechnology Information (NCBI) data bank using the following accession numbers: XM_028276483.1 (*DvvGr1*), XM_028280521.1 (*DvvGr2*), and XM_028272033.1 (*DvvGr3*). These gene sequences were translated to obtain protein sequences. The obtained protein sequences and the protein sequences of CO_2_ receptors from different insects were used to infer evolutionary relationships using the Neighbor-Joining method in MEGA 7 (***Kumar et al., 2016; Robertson and Kent, 2009; Rodrigues et al., 2016; Saitou and Nei, 1987***). The optimal tree with the sum of branch length = 4.44068889 is provided in Fig 1a. The percentage of replicate trees in which the associated taxa clustered together in the bootstrap test (100 replicates) are shown next to the branches (***Felsenstein, 1985***). The tree is drawn to scale, with branch lengths in the same units as those of the evolutionary distances used to infer the phylogenetic tree. The evolutionary distances were computed using the Poisson correction method (***Zuckerkandl and Pauling, 1965***) and are in the units of the number of amino acid substitutions per site. A total of 242 amino acid positions were included in the final dataset. Graphical representation and edition of the phylogenetic tree were performed with the Interactive Tree of Life (version 3.5.1)(***Letunic and Bork, 2016***). Protein tertiary structures and topologies were predicted using Phyre2 (***Kelley et al., 2015***).

### Production of double stranded RNA

*Escherichia coli* HT115 were transformed with recombinant L4440 plasmids that contained a 211-240 bp long gene fragment targeting one of the three CO_2_ receptors. Cloned nucleotide sequences were synthetized *de novo* (Eurofins, Germany). To induce the production of dsRNA, an overnight bacterial culture was used to inoculate fresh Luria-Berthani broth (25 g/L, Luria/Miller, Carl Roth GmbH, Karlsruhe, Germany). Once the bacterial culture reached an OD_600_ of 0.6-0.8, it was supplemented with isopropyl β-D-1-thiogalactopyranoside, IPTG) (Sigma-aldrich, Switzerland) at a final concentration of 2mM. Bacterial cultures were incubated at 37°C in an orbital shaker at 130 rpms for 16 additional hours. Bacteria were harvested by centrifugation and stored at −20°C for further use (***Kim et al., 2015***).

### Gene silencing experiments

To induce gene silencing in WCR, six to ten second instar WCR larvae were released in solo cups (30 mL, Frontier Scientific Services, Inc., Germany) containing approx. 2g of autoclaved soil (Selmaterra, Bigler Samen AG, Thun, Switzerland) and 2-3 freshly germinated maize seedlings. Maize seedlings were coated with 1 ml of bacterial solution containing approximately 200-500 ng of dsRNA, targeting the different CO_2_ receptor genes. As controls, larvae were fed with bacteria producing dsRNA targeting green fluorescent protein genes (GFP), which are absent in the WCR genome (***Rodrigues et al., 2016***). Fresh bacteria and seedlings were added to solo cups every other day for three consecutive times. Two days after the last dsRNA/bacteria application, larvae were collected and used for experiments.

### Gene expression measurements

Total RNA was isolated from approximately 10 mg of frozen, ground and homogenized WCR larval tissue (3-7 larvae per biological replicate, n=8) using the GenElute Universal Total RNA Purification Kit (Sigma-Aldrich, St. Louis, MO, USA). A NanoDrop spectrophotometer (ND-1000, Thermo Fisher Scientific, Waltham, MA, USA) was used to estimate RNA purity and quantity. DNase-treated RNA was used as template for reverse transcription and first strand cDNA synthesis with PrimeScript Reverse Transcriptase (Takara Bio Inc., Kusatsu, Japan). DNase treatment was carried out using the gDNA Eraser (Perfect Real Time) following manufacturer’s instructions (Takara Bio Inc.). For gene expression analysis, 2 μl of undiluted cDNA (i.e. the equivalent of 100 ng total RNA) served as template in a 20 μl qRT-PCR using the TB Green Premix Ex Taq II (Tli RNaseH Plus) kit (Takara Bio Inc.) and the Roche LightCycler 96 system (Roche, Basel, Switzerland), according to manufacturer’s instructions. Transcript abundances of the following WCR genes were analysed: *DvvGr1, DvvGr2, and DvvGr3* (***Rodrigues et al., 2016***). *Actin* was used as reference gene to normalize expression data across samples. Relative gene expression levels were calculated by the 2^-ΔΔCt^ method (***Livak and Schmittgen, 2001***). The following primers were used: DvvGr1-F CGTTAATTTAGCTGCTGTGG, DvvGr1-R GTTTTCTGTTGCTAGAGTTGC, DvvGr2-F GAACTAAGCGAGCTCCTCCA, DvvGr2-R CAGAAGCACCATGCAATACG, DvvGr3-F GCAACGCTTTCAGCTTTACC, DvvGr3-R GTGCATCGTCATTCATCCAG, DvvActin-F TCCAGGCTGTACTCTCCTTG, and DvvActin-R CAAGTCCAAACGAAGGATTG.

### Belowground olfactometer experiments with *DvvGr1*-,*DvvGr2*-,*DvvGr3*-silenced WCR larvae

To determine whether silencing putative CO_2_ receptor genes impairs the ability of WCR larvae to behaviourally respond to CO_2_, we silenced *DvvGr1*, *DvvGr2*, and *DvvGr3* as described above and conducted behavioural experiments using belowground olfactometers. Larvae that were fed bacteria expressing dsRNA targeting green fluorescent protein (GFP) were used as controls (herein referred as wild type larvae; WT). We conducted choice experiments using two-arm belowground olfactometers with carbonated water as a CO_2_ source (***Bernklau and Bjostad, 1998a; Huang et al., 2017; Jewett and Bjostad, 1996***). For this, a plastic cup containing 50 ml of carbonated water was placed in one arm of the olfactometer and a plastic cup containing 50 ml of distilled water was placed in the opposite arm (Fig. 2B). Five larvae were released in the middle of the olfactometer (indicated by a red arrow) and their positions were recorded 1h after their release. Seven olfactometers per larval type and experiment were assessed. The experiment was repeated twice. CO_2_ levels on each arm of the olfactometer were measured by gas chromatography coupled to a flame ionization detector (GC-FID) as described (***Zhang et al., 2019***).

### Insect behavioural experiments

To determine larval responses to plant volatile organic compounds other than CO_2_, we evaluated insect behaviour to methyl anthranilate. For this, we evaluated insect preferences for seedling roots or for seedling roots placed next to a filter paper disc treated with methyl anthranilate in choice experiments following a similar experimental procedure as described (***Bernklau et al., 2018a***). To this end, five 2^nd^-3^rd^ instar WCR larvae were released in the middle of a sand-filled petri plate (9 cm diameter, Greiner Bio-One GmbH, Frickenhausen, DE) where they encountered seedling roots in one side or seedlings roots and a filter paper disc treated with 10 μl of methyl anthranilate solution (10 mg/ml of water) in the other side. Ten Petri plates per larval type with five larvae each were evaluated (n=10). Methyl anthranilate was purchased from Sigma (CAS: 134-20-3; Sigma Aldrich Chemie, CHE). Larval preferences were recorded 1h after releasing the larvae.

To investigate larval responses to non-volatile root chemicals, we tested two well established attractants, Fe(III)(DIMBOA)_3_ and a sugar blend (***Bernklau et al., 2018b; Hu et al., 2018***). To evaluate the attractive effects of Fe(III)(DIMBOA)3, we released six 2^nd^-3^rd^ instar WCR larvae in the middle of a moist sand-filled Petri plate (6 cm diameter, Greiner Bio-One GmbH, Frickenhausen, DE) where they had a choice between a filter paper disc treated with 10 μl of Fe(III)(DIMBOA)_3_ (1 μg/ml of water) or, on the opposite side, a filter paper disc treated with water only. Twenty Petri plates with six larvae each were evaluated (n=20). Fe(III)(DIMBOA)3 was prepared fresh by mixing FeCl_3_ and DIMBOA at a 1:2 ratio as described. Larval preferences were recorded 1h after releasing the larvae. To evaluate attractive effects of soluble sugars for WCR larvae, we followed the procedure described by (***Bernklau et al., 2018b***). Briefly, we released six 2^nd^-3^rd^ instar WCR larvae in the middle of a moist sand-filled petri plate (6 cm diameter, Greiner Bio-One GmbH, Frickenhausen, DE) where they had a choice between a filter paper disc treated with 10 μl of a mixture of glucose, fructose, and sucrose (30 mg/ml of each sugar), or, on the opposite side, a filter paper treated with water only. Twenty Petri plates with six larvae each were assayed (n=20). Larval preferences were recorded 3h after release.

To evaluate the attractiveness of CO_2_ for WCR larvae, we conducted choice experiments using two-arm belowground olfactometers. CO_2_ levels were increased in one arm of the olfactometer by delivering CO_2_-enriched synthetic air (1% CO_2_, Carbagas, Switzerland). Olfactometer arms were closed on top using parafilm during CO_2_ delivery. A manometer connected to the synthetic air bottle allowed to fine-tune CO_2_ delivery rates. CO_2_ levels were measured using a gas analyzer (Li7000, Li-Cor Inc., Lincoln, Nebraska, USA). CO_2_ levels were increased at different levels from 22 to 1832 ppm above ambient CO_2_ levels. Once the desired CO_2_ concentrations were reached, CO_2_ delivery was terminated, olfactometer arms were connected to olfactometer central connectors and six 2^nd^-3^rd^ instar WCR larvae were release immediately, and their choice were evaluated within ten minutes of release. Three olfactometers per larval type were assayed (n=3).

To assess the impact of *DvvGr2* on larval motility, we followed the trajectories of individual larvae in Petri plate that were outfitted with a CO_2_ point releaser. For this, Petri plates (9 cm diameter, Greiner Bio-one, Austria) were pierced in their centers to allow the insertion of a needle that released CO_2_ at 60 ppm above ambient level. Larvae were released at the rim of the Petri plates (4.5 cm apart from CO_2_ source). Petri plates were overlaid with moist filter paper (9cm diameter, GE Healthcare, UK). Insect behavior was observed for three minutes. Six Petri plates with one larva each were assayed (n=6). In a second experiment, we followed the trajectories of individual larvae in Petri plate with maize roots. For this, two root pieces (3-4 cm long) of four-day-old maize seedlings were placed at the rim of a Petri plate overlaid with moist filter paper. Then, on the opposite rim (9cm apart), we released one 2^nd^-3^rd^ instar WCR larva and followed their trajectories during three minutes. Six Petri plates with one larva each were assayed (n=6). Crawled distances were measured using ImageJ 1.53a.

### Host location experiments

To determine the importance of plant-derived CO_2_ for distance-dependent host-location by WCR larvae, we used two experimental setups: two-arm belowground olfactometers and soil-filled trays (Fig. 5, Fig. 6). For behavioural experiments using belowground olfactometers, we used two types of olfactometers: one with short arms that allowed for the release of larvae at 9 cm (Fig. 5A) from plant volatile sources, and one with long arms that allow to release the larvae at 18 cm (Fig. 5B) from plant volatiles sources. To specifically investigate the importance of plant derived CO_2_, we contrasted insect behavioural responses to intact plant odours and to plant odours without CO_2_. CO_2_ was experimentally removed using soda lime (Carl Roth, Karlsruhe, Germany). For this, a layer of 5g of soda lime granules (2-4mm) were placed between Teflon connectors and sand 30 min before the experiments. Olfactometers without soda lime served as controls. Olfactometer arms were filled with sand 48h before the experiment and one three-week-old maize plant was transplanted to one arm of the olfactometer. Four olfactometers with six larvae each were assayed (n=4). CO_2_ levels were measured using a gas analyzer (Li7000, Li-Cor Inc., Lincoln, Nebraska, USA). For behavioural experiments using soil-filled trays, second instar WCR larvae were released at 16, 32, 48, or 64 cm from four or five 2-to 3-week-old maize plants. Larval release points are indicated by colour arrows (Fig. 6B). Plants were planted in plastic trays (80cmx15cmx4.5cm)(Migros do it + garden, Switzerland) filled with soil. Eight h after releasing the larvae, their positions were recorded. Six trays per larval type and distance with twenty larvae each were assayed (n=6). CO_2_ levels in the soil gas phase of each tray zone were measured by GC-FID as described (***Zhang et al., 2019***).

### Host finding behavioural experiments with nutrient-deficient plants

Optimally fertilized plants emit higher levels of respiratory CO_2_ than sub-optimally fertilized plants (***Zhu and Lynch, 2004***). To test whether WCR larvae orients towards and prefer optimally fertilized maize plants and to evaluate the importance of CO_2_ detection through *DvvGr2* in this context, we silenced this gene and conducted choice experiments to contrast larval preference for plants that were differentially fertilized. To this end, we fertilized maize plants with three doses of fertilizer: 0.1% (optimally fertilized), 0.05% (medium fertilized), or 0.01% (low fertilized). Plantaaktiv^®^ 16+6+26 Typ K fertilizer (Hauert HBG Dünger AG, Grossaffoltern, Switzerland) was dissolved to the abovementioned concentrations and applied following manufacture’s indications. Fertilizer macro- and micronutrient composition are described above (Plants and planting conditions). For the choice experiments, plants were grown under the different fertilizer regimes. 24 hr before the choice experiment, five 15-day-old, optimally fertilized plants were transferred and re-planted in one corner of an 80 cm long plastic tray (Migros do it + garden, Switzerland) filled with soil (Figs 5a, b). At the opposite side, plants grown either under low or medium fertilizer regimes were equally transferred and re-planted (Figs 7A, B). Then, twenty 2^th^ instar WCR larvae were released in the middle of the tray (Zone 3, Fig 6A, B, red arrows). Six independent trays per larval type and fertilizer regime pair were evaluated (n=6). Eight hours after releasing the larvae, larval positions were recorded.

### Effect of plant nutritional status on larval growth

To test whether larval preference for optimally fertilized plants is reflected in their growth, we measured larval weights of larvae feeding on roots of plants that were grown under the different fertilizer regimes. Fertilizer regimens are described above. For this, seven 1^th^ instar larvae were released into solo cups (30 mL, Frontier Scientific Services, Inc., DE) containing approx. 2g of organic soil (Selmaterra, Bigler Samen AG, Thun, CHE). Fresh roots of 15-day old plants were provided everyday *ad libitum*. Roots were washed thoroughly to remove fertilizer traces from the root surface. Twenty solo cups per treatment were included (n=20). Eight days after the beginning of the experiment, larvae were weighed using a micro balance. Four to seven larvae were recovered per experimental unit at the end of the experiment.

### Statistical analyses

Differences in gene expression levels, carbon dioxide concentrations, and larval performance were analysed by analysis of variance (ANOVA) using Sigma Plot 12.0 (SystatSoftware Inc., San Jose, CA, USA). Normality and equality of variance were verified using Shapiro–Wilk and Levene’s tests, respectively. Holm–Sidak post hoc tests were used for multiple comparisons. Datasets from experiments that did not fulfill the assumptions for ANOVA were natural log-, root square-, or rank-transformed before analysis. The differences in larval preference were assessed using Generalized Linear Models (GLM) under binomial distribution and corrected for overdispersion with quasi-binomial function when necessary followed by analysis of variance (ANOVA) and FDR-corrected post hoc tests. All analyses were followed by residual analysis to verify the suitability of the error distribution and model fitting. All The above analyses were conducted using R 3.2.2 (43) using the packages “lme4”, “car”, “lsmeans” and “RVAideMemoire” (***Bates et al., 2014; Fox and Weisberg, 2011; Herve, 2015; Lenth, 2016; Team, 2014***).

## Acknowledgements

We thank Evangelia Vogiatzaki and Celine Terrettaz for technical assistance, the members of the Research Section Biotic Interactions of the Institute of Plant Sciences of the University of Bern, Switzerland for their support and helpful discussions. This project was supported by the University of Bern, a European Union Horizon 2020 Marie Sklodowska-Curie Action (MSCA) Individual Fellowship (Grant Nr. 794947 to B.C.J.S.), and the Swiss National Science Foundation (Grant Nr. 155781 to M.E., Grant Nr. 186094 to R.A.R.M).

## Author contributions

Conceptualization, M.E. and R.A.R.M.; Methodology, R.A.R.M. and M.E.; Formal analysis, R.A.R.M., M.E., and C.C.M.A.; Investigation, R.A.R.M., C.C.M.A., V.T., B.C.J.S., and G.J.; Visualization, R.A.R.M.; Supervision, R.A.R.M. and M.E.; Funding Acquisition, M.E., and R.A.R.M Writing – original draft, R.A.R.M. and M.E.; Writing – review & editing, R.A.R.M., M.E., B.C.J.S., and C.C.M.A.

## Data availability

All data sets are available from Dryad (to be added at a later stage).

## Supplementary Figures

**Figure S1.**
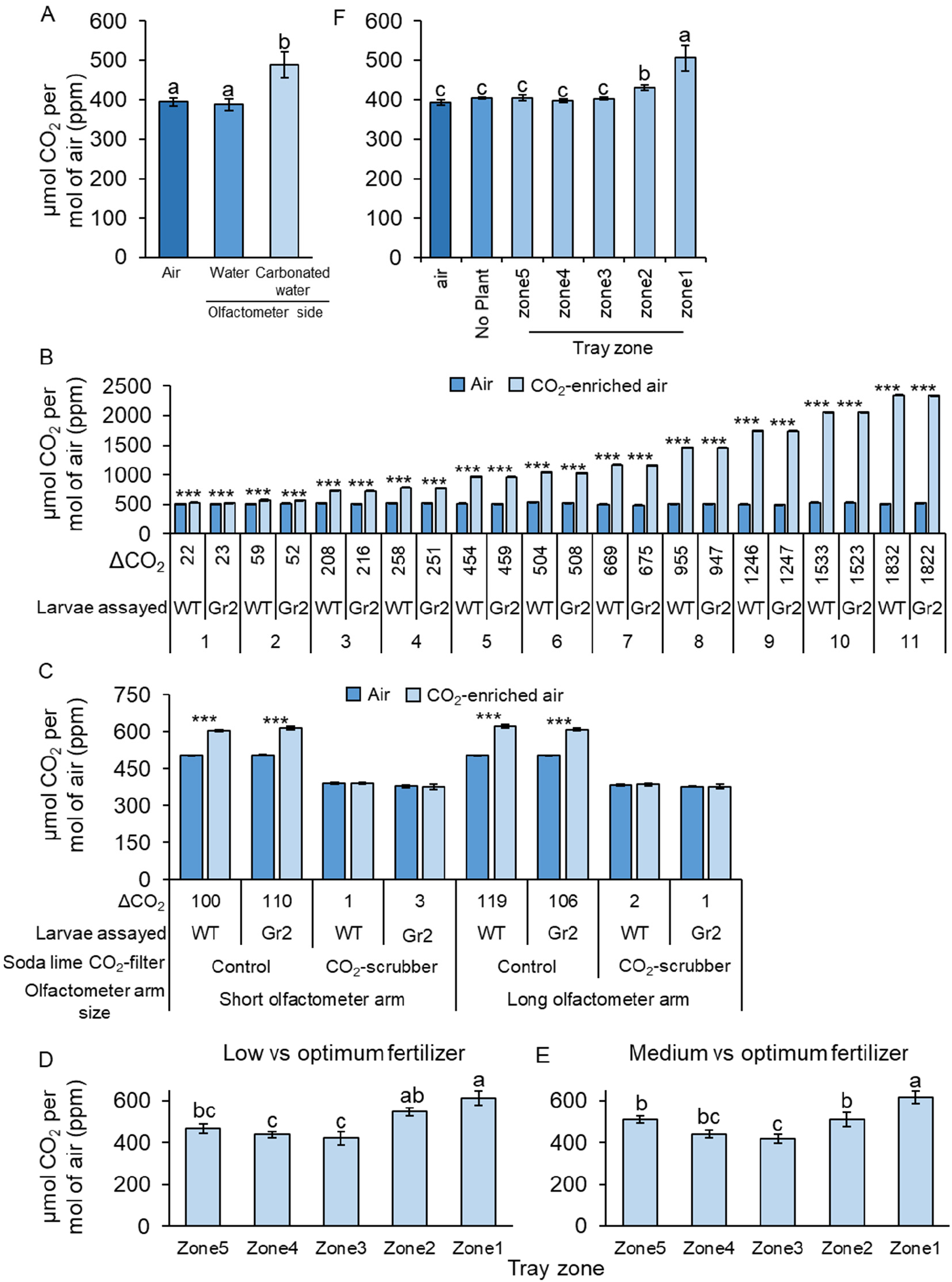
Carbon dioxide measurements for the different experiments. Mean (±SEM) CO_2_ levels measured in behavioural experiments of Figure 2 (A), Figure 3 (B), Figure 5 (C), Figure 7 (D, E), Figure 6 (F). Different letters indicate significant differences in CO_2_ levels (*p*<0.05 by one-way ANOVA with Holm’s multiple-comparisons test). Asterisks indicate significant differences in CO_2_ levels (***: *p*<0.001 by ANOVA with Holm’s multiple-comparisons test). Gr2: *DvvGr2*-silenced larvae. For details on CO_2_ sampling points, refer to the main figures.

**Figure S2.**
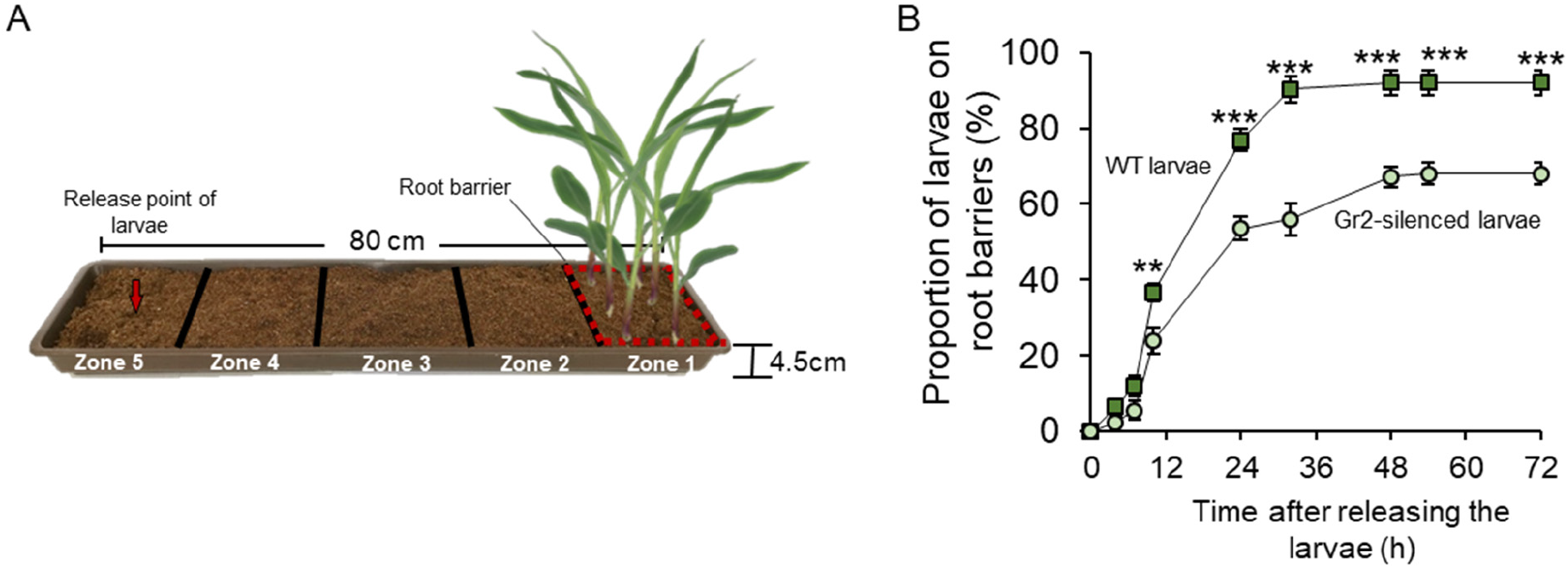
Root-derived CO_2_ is required for host location by WCR larvae at long distances. (A) Schematic representation (photomontage) of experimental set up used. (B) Mean (±SEM) proportion of WCR larvae retrieved at close vicinity of root systems (i.e.: on root barriers). Twenty-five larvae were released at 64cm from the plants and five arenas were assayed (n=5). Soil surrounding root barriers (indicated by dashed red lines) were carefully inspected and the observed larvae were collected and counted regularly. Asterisks indicate statistically significant differences in the proportion of larvae retrieved at each time point (**: *p*<0.01; ***: *p*<0.001 by two-way repeated measures ANOVA with Holm’s multiple comparisions test). For *P*-values of the different factors and their interactions, refer to Table S1.

